# Identification of sex chromosomes using genomic and cytogenetic methods in a range-expanding spider, *Argiope bruennichi* (Araneae: Araneidae)

**DOI:** 10.1101/2021.10.06.463373

**Authors:** Monica M. Sheffer, Mathilde Cordellier, Martin Forman, Malte Grewoldt, Katharina Hoffmann, Corinna Jensen, Matěj Kotz, Jiří Král, Andreas W. Kuss, Eva Líznarová, Gabriele Uhl

**Author notes:** Correspondence address: Monica M. Sheffer, Zoological Institute and Museum, University of Greifswald, Loitzer Strasse 26, 17489 Greifswald, Germany Correspondence. Department of Biology, University of Washington, 98195 Seattle, WA, USA. Department of Molecular Biology and Genetics, Aarhus University, 8000 Aarhus, Denmark (Malte Grewoldt).

## Abstract

Differences between sexes in growth, ecology and behavior strongly shape species biology. In some animal groups, such as spiders, it is difficult or impossible to identify the sex of juveniles based on external morphology. This information would be useful for field surveys, behavioral experiments, and ecological studies on e.g. sex ratios and dispersal. In species with sex chromosomes, sex can be determined based on the specific sex chromosome complement. Additionally, information on the sequence of sex chromosomes provides the basis for studying sex chromosome evolution. We combined cytogenetic and genomic data to identify the sex chromosomes in the sexually dimorphic spider *Argiope bruennichi*, and designed RT-qPCR sex markers. We found that genome size and GC content of this spider falls into the range reported for the majority of araneids. The male karyotype is formed by 24 acrocentric chromosomes with an X_1_X_2_0 sex chromosome system, with little similarity between X chromosomes, suggesting origin of these chromosomes by X chromosome fission or early duplication of an X chromosome and subsequent independent differentiation of the copies. Our data suggest similarly sized X chromosomes in *A. bruennichi*. They are smaller chromosomes of the complement. Our findings open the door to new directions in spider evolutionary and ecological research.

## Introduction

In most animals, sex is genetically determined. Genotypic mechanisms of sex determination are very diverse, depending, for example, on the ratio of X chromosomes to autosomes, on sex-determining factor(s) localized on autosomes and/or sex chromosomes, on the presence or absence of a sex chromosome specific for the heterogametic sex (Y or W), or on ploidy level as in many Hymenoptera (Bachtrog *et al*., 2014; Hamm, Meisel, & Scott, 2015; Vicoso, 2019).

In animal species with genotypic sex determination, females and males often differ in the type and/or number of sex chromosomes, with many variations. Knowledge on the karyotype and the specifics of the sex chromosomes provides the basis for studying a plethora of ecological and evolutionary questions. For ecological studies, it is desirable to know the sex of collected individuals. Offspring sex allocation, sex-specific growth and dispersal strategies, and sex differences in early developmental pathways and physiological plasticity can then be investigated (Cordellier *et al*., 2020). However, in many animal taxa, the sexes cannot easily be identified in adults and even less so in juveniles. The sex of early stages is often only determinable based on their internal anatomy (presence of testis or ovaries), or on their sex chromosome complement (e.g., Avilés & Maddison, 1991).

Spiders belong to one of the most diverse animal orders (Coddington & Levi, 1991). While they play an important role in terrestrial ecosystems, it is impossible to sex juvenile spiders in the field. Spiders possess complex sex chromosome systems. To date, karyotypes of more than 800 spider species have been characterized (Araujo *et al*., 2021). In 67% of karyotyped spiders (calculated at the time of publication), and particularly in the entelegyne spiders, which are the most studied, there are two X chromosomes, X_1_ and X_2_ (X_1_X_2_0 system), where the male exhibits one copy and the female two copies (_⍰_X_1_X_2_/⍰X_1_X_1_X_2_X_2_) (Araujo *et al*., 2012). This pattern is probably ancestral for entelegynes (Král *et al*., 2006). Systems with three X chromosomes or more also exist. These X chromosomes are supposedly non-homologous based on their achiasmatic pairing during male meiosis (Kořínková & Král, 2013). Some lineages of entelegyne spiders exhibit an X0 system, which has originated by X chromosome fusions, or neo-sex chromosomes, which arose by rearrangements between the X chromosome(s) and autosome(s). Neo-sex chromosome systems contain both X and Y chromosome(s). Systems formed by two or more X chromosomes are termed multiple X chromosome systems (White, 1973).

Due to the different number of X chromosome copies in males and females, sexing could be achieved in spiders once targeted molecular markers for X chromosomes are available: the relative copy number of sequences on the X chromosomes, with males having half the number of copies relative to females, could be assessed through quantitative PCR. Sexing performed in this manner can be done at a high throughput, making it reasonable for ecological studies with relatively large sample sizes.

In recent years, several approaches to identify X chromosomal sequences have been introduced that rely on high throughput sequencing methods. These methods were reviewed by Palmer *et al*. (2019), and vary in their costs, required sample sizes, and on the availability of a reference genome (Al-Dous *et al*., 2011; Picq *et al*., 2014; Gautier, 2014; Hou *et al*., 2015; Muyle *et al*., 2016). In species with a chromosome-level genome assembly, comparing the sequencing coverage depth across chromosomes between individuals of each sex is a reliable method for identifying sex chromosomes. Females, as the homogametic sex, would show twice as much coverage as males on X chromosomes (i.e. (Vicoso & Bachtrog, 2011; Fraïsse, Picard, & Vicoso, 2017).

In 2021, Sheffer *et al*. published an annotated genome assembled to haploid chromosome level of an entelegyne orb-weaving spider, *Argiope bruennichi* (Scopoli, 1772). With this chromosome-level genome, we can employ the coverage depth approach to identify sex chromosome sequences in this species. *A. bruennichi* is a sexually dimorphic species with a Palearctic distribution, which has rapidly expanded its range in Europe over the course of the last century (Krehenwinkel & Tautz, 2013; Krehenwinkel, Rödder, & Tautz, 2015; Wawer *et al*., 2017). Sex differences in early life stages of these spiders are particularly interesting since juveniles perform aerial dispersal with silk (ballooning), a behavior which is restricted to the first few instars (Krüger, 2014) and might be sex biased (Krehenwinkel *et al*., 2016).

The *A. bruennichi* genome assembly was based upon female specimens, and is comprised of 13 scaffolds, which suggests 2n = 26. This chromosome number is found in females of most orb-weaving spiders, and it is consistent with the diploid chromosome number of the species published by Zhang & Tong (1990). This study, however, contains no information on the number or sex of the individual(s) examined. With additional karyological and sequence data for males, it would be possible to ascertain the karyotype/sex determination system of the species and identify the sex chromosomes in the published assembly.

We set out to (i) determine the karyotype and genome size of *A. bruennichi*, (ii) identify which superscaffolds represent sex chromosomes in the genome assembly, and (iii) develop molecular markers for sexing spiders using RT-qPCR.

## Methods

### Chromosome preparations and their evaluation

We collected eight subadult *A. bruennichi* males for chromosomal analysis at village Březová, Czech Republic (49.901N, 13.887E) in late June 2021. The number of meiotic cells in the testes of these males was lower than found in subadult or young adult males of other araneids (Kotz and Forman, unpublished observation). We followed the protocol of Dolejš *et al*. (2011) for preparation of chromosome slides and Giemsa staining. We inspected the slides under an Olympus BX 50 microscope and photographed selected plates using an Olympus DP 71 CCD camera. We measured chromosome lengths using ImageJ (https://imagej.nih.gov/ij/index.html). The length ratio of sex chromosomes was established using 20 randomly selected diakinesis and metaphase I plates. Chromosome morphology was based on centromere position, which was apparent during early metaphase II.

### Determination of genome size and GC content

We determined the diploid genome size (2C value) and guanosine cytosine (GC) content of *A. bruennichi* by flow cytometry (FCM) using five adult males and five females (collected in July 2020, Albrechtice, Czech Republic, 49.924N, 16.658E, stored at −80IZ°C). Sample preparation was based on the protocol of Král *et al*. (2019) with a different buffer. We chopped the legs together with an internal plant standard in GBP buffer (Loureiro *et al*., 2007), with the pH adjusted to 9.5 and 1.5% (v/v) of Triton X-100 (Sigma-Aldrich) and 1.5% (v/v) of polyvinylpyrrolidone (Sigma-Aldrich). After filtration through a nylon sifter, we added fluorescent dye. We performed two concurrent FCM gauges, using: (i) Propidium iodide (PI, 50 μg/ml) – a nonspecific dye to establish genome size of the sample – and (ii) 4⍰,6-Diamidine-2⍰-phenylindole dihydrochloride (DAPI, 4 μg/ml), a fluorochrome with higher specificity to AT pairs. The resulting histograms were evaluated using FloMax software (Partec). We calculated the 2C value as the ratio of PI fluorescence of sample and standard. We compared PI and DAPI ratios to establish GC content as described in šmarda *et al*. (2008). We measured each individual twice for PI and twice for DAPI to account for machine fluctuations. We used fresh leaves of *Pisum sativum* cultivar ‘Ctirad’ as an internal standard. We used the following genomic values of this plant for calculations: 2C=7841.27 Mbp, GC=41.77% (Veselý *et al*., 2012). Primary data used for calculation of 2C and base content are available in Supplementary Table S1.

### Whole-genome sequencing of male and female Argiope bruennichi

#### DNA extraction and library preparation

We will refer to chromosome-level scaffolds in the genome assembly as “superscaffolds” and number all of them by length. For karyotype, we use the traditional approach and number autosomes by length and refer to X chromosomes separately.

In order to identify the X chromosomes in the *A. bruennichi* genome assembly, we sequenced one adult male and one adult female collected near Klausdorf, Germany at the end of July 2020 (54.424N, 13.029E). We extracted DNA from eight legs of the male and four legs of the female, as follows: we disrupted the leg tissue with a mortar, pestle, and liquid nitrogen. The powdered tissue was transferred into tubes containing Proteinase K and cell lysis buffer (10mM Tris pH 8, 100mM NaCl, 10mM EDTA pH 8, 0.5% SDS, double-distilled water). We incubated the tubes overnight at 55°C, cooled them to room temperature and added RNAse A. We used 5M NaCl to precipitate proteins and Isopropanol to precipitate DNA. We cleaned the DNA extract using 70% Ethanol and eluted it in TE buffer. We prepared the libraries with the NEBNext Ultra II FS DNA Library Prep Kit (New England Biolabs, Ipswich, Massachusetts, USA) according to the manufacturer’s protocol. We indexed and amplified the libraries with six PCR cycles, then pooled them and sequenced the 2×75 bp paired-end libraries on one Flowcell of an Illumina NextSeq 550 with a High Output v2 kit (150 cycles) (Illumina Inc., San Diego, California, USA).

#### Bioinformatic processing and coverage comparison

The reads were demultiplexed using the Illumina bcl2fastq tool in BaseSpace. We removed adapter sequences using AdapterRemoval (v.2.2.2) (Schubert, Lindgreen, & Orlando, 2016) with the --identify-adapters flag. Read quality was assessed using FastQC (v0.11.8) (Andrews, 2010); all reads were high quality, and no further quality trimming was necessary. Adapter-trimmed reads are available under BioProject PRJNA629526. We mapped the reads of each specimen onto the *A. bruennichi* genome assembly using BWA-MEM (v.0.7.12-r1039) with default settings (Li, 2013), sorted and indexed the mapped reads using SAMtools sort and SAMtools index (SAMtools v.1.3.1) (Li *et al*., 2009), then calculated the coverage of each of the 13 superscaffolds using QualiMap (v.2.2.1) (Okonechnikov, Conesa, & García-Alcalde, 2016).

### Development of sex markers using RT-qPCR

#### DNA extraction

In total, we extracted genomic DNA from six males collected in Greifswald (54.093N, 13.366E) and six females collected near Klausdorf (54.424N, 13.029E), Germany at the beginning of August 2020, following either the procedure outlined above, or a protocol based on the Promega ReliaPrep™ gDNA Tissue Miniprep System (see Table S2). Prior to using the Promega kit, we ground two legs each from females or four legs each from males with a Dounce homogenizer; we added 200 μl of Tail Lysis Buffer and 30 μl of Proteinase K Solution to the sample, and extracted the DNA following the manufacturer’s instructions. We carried out the final elution step twice to a total elution volume of 100 μl.

Further, we extracted DNA from ten first instar spiderlings that came from one egg sac, laid by a female collected in Pärnu, Estonia in 2018 (58.297N, 24.597E). Genomic DNA was extracted as described in the preceding section. Whole spiderling bodies were used, rather than legs, due to their small size. First instar spiderlings from this population have an average mass of approximately 0.3 mg (Sheffer, personal observation). We determined the DNA concentration for each sample using a NanoDrop spectrophotometer (Thermo Scientific, Wilmington, DE) and a Qubit 4 Fluorometer with the Quant-iT™ Qubit™ dsDNA BR Assay Kit (Thermo Scientific, Wilmington, DE). DNA concentrations can be found in Table S2.

#### Primer design, PCR and RT-qPCR

We designed primers for loci on the putative X chromosomes (superscaffolds 9 and 10 in the *A. bruennichi* assembly; see results section) and 2 putative autosomes, superscaffolds 5 and 7, using Primer3 (version 4.1.0; (Untergasser *et al*., 2012) with the following parameters: length between 20 and 25 nucleotides, melting temperature between 57 and 62°C, region amplified between 150 and 200 bp in length, with one GC clamp (Table S3). We used additional data to select the loci, such as the location of homologous sequences in spiders *Trichonephila senegalensis* (Araneidae) (Walckenaer, 1841) (Grewoldt *et al*., unpublished data) and *Stegodyphus mimosarum* (Eresidae) (Pavesi, 1883) (Bechsgaard *et al*., 2019). We tested the amplification with conventional PCR, checked on a 1.5% agarose gel with ethidium bromide staining and UV transillumination. We then conducted RT-qPCR using the EvaGreen® qPCR-Mix II, on a ThermoFisher StepOne™ Real-Time PCR System (Thermo Fisher Scientific, Waltham, USA). Primer efficiencies were measured by diluting 20 ng/μl of template DNA from a female in a stepwise manner, 1:10 down to the lowest concentration of 0.02 ng/μl, to obtain a standard curve. After ensuring comparable efficiencies for all primer pairs, we used DNA from the six females and males as templates and simultaneously measured amplification in real time for each primer pair in triplicate. One non template control was used for each primer combination. All amplification protocols were conducted with the following reaction mix: 2 μL EvaGreen® qPCR-Mix II 5x, 0.2 μL forward primer, 0.2 μL reverse primer, 1μL template DNA, and 6.6 μL nuclease free water. We used the following program: initial denaturation for 15 minutes at 95°C, 40 cycles of denaturation (15 seconds at 95°C), annealing (20 seconds at 60°C), and extension (20 seconds at 72°C), with a final extension at 72°C for 10 minutes, and a melting curve (ramping from 65°C to 95°C). Ct values were collected for further analysis (Table S2).

After validation of the method and determination of effective markers (see results) using the adult specimens, we used a subset of markers (see Tables S2 and S3) to determine the sex of the spiderlings.

#### RT-qPCR data analysis

We applied the 2^-ΔΔCt^ method according to (Rao *et al*., 2013) to analyze the RT-qPCR data. To account for different primer efficiencies (E_real_), the Ct values were normalized to correspond to an optimal efficiency of 100% (E_opt_). We averaged the Ct values of the three technical replicates; if any of the three replicates showed a standard deviation >1, it was excluded from the average. The ΔCt value was calculated from the difference between reference sequences (autosomal loci) and a target sequence of interest (X-linked locus). Further, the female data points were used as the reference, as females possess two sex-linked and two autosomal gene copies. Consequently, if the number of gene copies between females and males is the same, a value of 1 is expected, whereas if the number of gene copies in males is half that in females, a value of 0.5 is expected.

## Results

### Chromosomes

The male karyotype of *A. bruennichi* was formed by 24 acrocentric chromosomes; the size difference between chromosomes was gradual from the longest to the shortest (Figure 1A). Analysis of male meiotic division revealed an X_1_X_2_0 sex chromosome system (Figure 1B, E-J). Chromosomes X_1_ and X_2_ were similar in length; X_2_ length corresponded to 91.3% (±5.3) of X_1_. We observed nearly the complete course of male meiotic division except anaphase I, telophase I, and telophase II. X chromosomes were positively heteropycnotic (i.e., they stained more intensively) from the beginning of meiosis until diakinesis (Figure 1B-E). The intensity of heteropycnosis decreased during metaphase I; X chromosomes were isopycnotic or slightly positively heteropycnotic during this period (Figure 1F). Positive heteropycnosis reappeared during prophase II (Figure 1G). During metaphase II (Fig. 1H), X chromosomes exhibited a slight positive heteropycnosis only. This pattern was also displayed at late anaphase II (Figure 1J). Tight pairing of X_1_ and X_2_ chromosomes was initiated at zygotene (Figure 1C). During zygotene and pachytene, sex chromosomes formed a body on the periphery of the plate (Figure 1C, D). During late prophase I (diplotene-diakinesis), X chromosome pairing was restricted to centromeric regions (Figure 1E). From metaphase I onwards, X chromosomes were arranged in parallel (Figure 1F, G, J).

**Figure 1.**
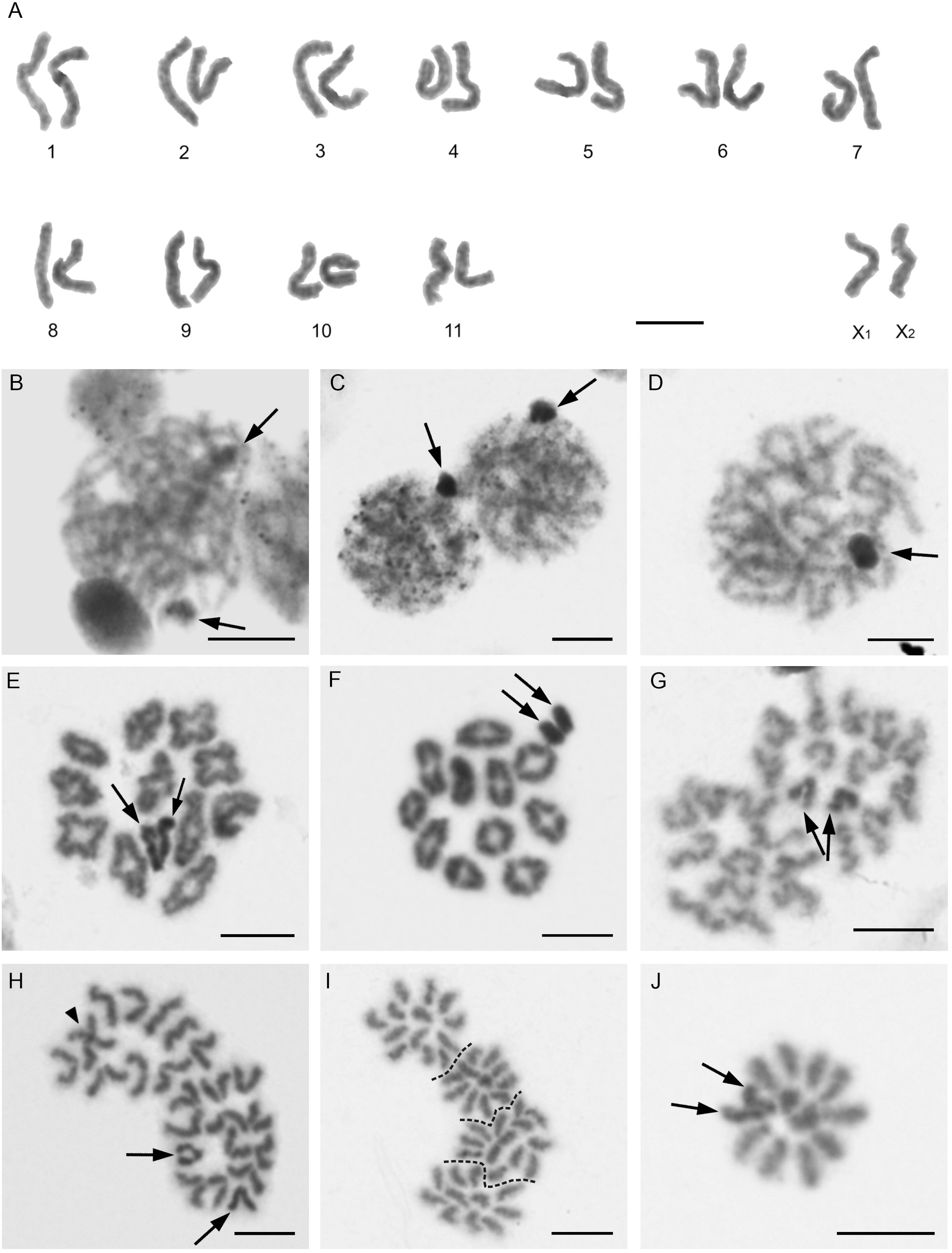
*A. bruennichi male*, 2n=24, X_1_X_2_0, Giemsa stained mitotic (A) or meiotic (B-J) chromosomes. Arrows points to sex chromosomes. (A) Karyotype, based on spermatogonial prometaphase, (B) leptotene. Chromosomes X_1_ and X_2_ are not associated, (C) two zygotene plates. Sex chromosomes form the body on the periphery of the nucleus, (D) pachytene. The sex chromosome body persists on the periphery of the nucleus, (E) diplotene. Heteropycnotic X_1_ and X_2_ pair by their centromeric regions, (F) metaphase I. Note a slight positive heteropycnosis of X chromosomes, which are arranged in parallel on the periphery of the plate, (G) two sister prophases II. The left plate contains 11 chromosomes and the right plate 13 chromosomes, (H) two sister metaphases II. Sex chromosomes are slightly positively heteropycnotic (arrowhead - two overlapping chromosomes), (I) two sister anaphases II. Two half plates with 11 chromosomes and two half plates with 13 chromosomes (half plates separated by dashed line). X chromosomes indistinguishable from the other chromosomes, (J) late anaphase II, half plate. Note a slight positive heteropycnosis of X chromosomes. Scale bar = 5 μm.

### Genome size and GC content

Female 2C was 4,079.809 Mbp (standard error (SE) ±256.265) and male 2C 3,921.083 Mbp (SE ±207.799). The average chromosome size (2C/2n) was 160.147 Mbp (156.916 Mbp based on female and 163.378 Mbp based on male data, respectively). X chromosomes contained 158.727 Mbp of DNA, based on the difference in female and male 2C. Taking in account the length ratio of X chromosomes, size of X_1_ chromosome can be estimated as 82.973 Mbp and size of X_2_ as 75.754 Mbp. The size of the X chromosomes was below the average chromosome size. Therefore, X chromosomes belonged to small chromosomes of the karyotype. The genome contained a low GC proportion, 35.500 % of GC (SE ±1.289) in females and 36.730% (SE ±1.189) in males, respectively.

### Coverage of male and female

The mean coverage (± standard deviation) across the 13 superscaffolds was 15.34 ±2.57 fold for the male and 16.50 ±0.76 fold for the female. We divided the coverage of the male for each superscaffold by the coverage of the female, to get the male’s relative coverage. 11/13 superscaffolds have nearly equal coverage of male relative to female, while two superscaffolds (9 and 10) have half coverage in the male (Figure 2A).

**Figure 2.**
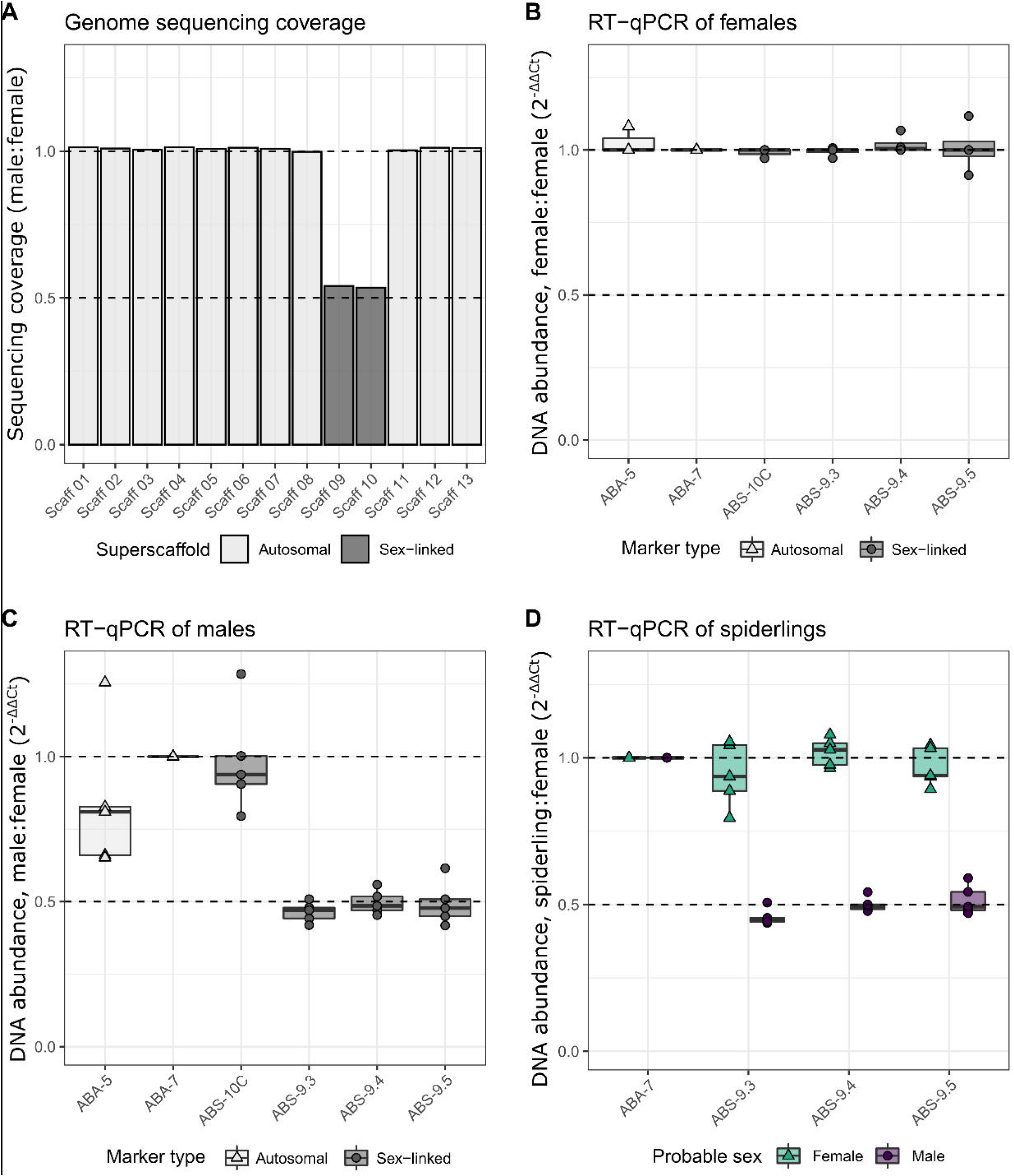
Identification of the sex chromosomes in the *Argiope bruennichi* genome assembly. The dashed horizontal lines represent (A) the expected coverage and (B-D) expected DNA abundance for sex chromosomes in females (1) and males (0.5) relative to females. (A) Relative sequencing coverage of males and females across the 13 superscaffolds (Scaff) in the genome assembly. (B-D) Boxplots show the results of the 2^-(ΔΔCt)^ method, calculated from average adjusted Ct values for each locus and individual. Symbols represent individual datapoints, boxes represent the interquartile range, thick lines represent medians, and whiskers represent 1.5 times the interquartile range. (B) RT-qPCR results for females. (C) RT-qPCR results for males. (D) RT-qPCR results for spiderlings for the subset of markers delivering reliable results (ABS-9.3, ABS-9.4, ABS 9.5, ABA-7).

### RT-qPCR

#### Sex marker validation in adults

To compare sex-linked with autosomal loci, the efficiency of the primers for each locus were evaluated. Primer efficiencies ranged from 83.3-147%, for the chosen loci (Table S3). Likewise, a melting curve was generated for each locus, confirming specific amplification during RT-qPCR. X-linkage was validated using RT-qPCR on DNA extracted from six females and six males (Figure 2B, C). The respective loci, sex-linked as well as autosomal, were normalized against each autosomal locus, as well as the average of all autosomal loci used. After comparing these normalization methods, we chose to normalize all loci against one autosomal locus, ABA-7, as this provided consistent results during validation. To validate the markers on individuals with known sex, the relative DNA abundance, or fold-change, in adult males and females, in comparison to females, was determined using the 2^-(ΔΔCt)^ method (Figure 2B, C). For females, autosomal loci showed an average fold-change of 1.09, and sex-linked loci showed an average fold-change of 1.00. One male individual was excluded from our dataset due to inexplicable fold-change values. With this male excluded, for a total of five males, the sex-linked loci on superscaffold 9 exhibited on average a 0.42-0.61 fold-change in DNA abundance in males, while the locus on superscaffold 10 had an average fold-change of 0.99 in males. The presumably autosomal locus on superscaffold 5 had an average fold-change of 0.84 in males (Figure 2C). Sex-linked loci on superscaffold 9 thus show about half the DNA abundance in males with an average of 0.48 compared to females, and an average of 1.01 for females compared to females. These markers on superscaffold 9 are therefore reliable for determining the sex of juveniles. Due to their inconsistent results, ABA-5 and ABS-10C were not used for sexing juveniles.

#### Application of sex markers in juveniles

We used one autosomal locus (ABA-7) and three sex-linked loci (ABS-9.3, ABS-9.4, and ABS-9.5) to determine the probable sex of ten individual spiderlings from a single clutch (Table S4). As shown in Figure 2D, two clear groups of spiderlings are visible: five individuals have an average fold change of 0.49 for the sex-linked loci, indicating that they are males, while the other 5 had an average fold change of 0.98 for these loci, indicating females. In this clutch, our newly-developed markers thus indicate an unbiased sex-ratio of 1:1.

## Discussion

Our study determined the male karyotype of *A. bruennichi*. It is composed of 24 chromosomes with an X_1_X_2_0 system. The same karyotype was found in two other *Argiope* species, namely *A. minuta* (Datta & Chatterjee, 1988), and *A. pulchella* (Bole-Gowda, 1958). This karyotype is the most common in analyzed species of the superfamily Araneoidea, and is probably an ancestral feature of this clade (Araujo *et al*., 2015). As in most other entelegyne spiders, the karyotype of *A. bruennichi* is formed by acrocentric chromosomes (Kořínková & Král, 2013); this karyotype structure can be considered ancestral for entelegynes (Král *et al*., 2006). Data on three other *Argiope* species are incomplete or different from *A. bruennichi*: the male karyotype of *A. amoena* is also formed by 24 chromosomes, including the X_1_X_2_0 system, but the chromosome morphology was not determined (Suzuki, 1951). The same diploid number was found in *A. catenulata* (Amalin, Barrion, & Rueda, 1992) and *A. luzona* (Carandang & Barrion, 1994), but the chromosome complement of *A. luzona* is reported to consist of both monoarmed (acrocentric and subtelocentric) and biarmed (metacentric and submetacentric) chromosomes (Carandang & Barrion, 1994). This karyotype structure is exceptional in entelegynes, and should be revisited by analysis of metaphase II, when centromeres are unequivocally identifiable. Biarmed chromosomes of *A. luzona* could arise from acrocentric chromosomes, such as those found in *A. bruennichi*, by pericentric inversions. The karyotype of *A. catenulata* is reported to be composed of holocentric chromosomes with an XY system (Amalin *et al*., 1992). In this case, the authors have most likely mistaken acrocentric chromosomes for holocentric ones, as they do not report any features which support holocentric structure of chromosomes. Moreover, other studied *Argiope* species have monocentric chromosomes (Kořínková & Král, 2013). Furthermore, the determination of the XY system by Amalin *et al*. (1992) is probably also incorrect, as they do not provide any supporting information on the mode of sex chromosome pairing or sex chromosome segregation during male meiosis.

As mentioned above, our karyotype analysis revealed 2n=24, X_1_X_2_0 in male *A. bruennichi*. Given the X_1_X_2_0 system, where males have one copy each of the two X chromosomes, this result fits with the 13 superscaffolds in the haploid genome assembly of a female *A. bruennichi* published by Sheffer *et al*. (2021): 11 autosomes + X_1_ + X_2_. The result showing half coverage for superscaffolds 9 and 10 in the male indicates that these represent the X chromosomes in the *A. bruennichi* genome assembly. These superscaffolds are very similar in length (123.24 Mbp and 122.82 Mbp), which does not match the lengths of X_1_ and X_2_ (82.973 Mbp and 75.754 Mbp) derived from the comparison of male and female genome sizes, with the X_1_:X_2_ ratio from image-based measurements. However, the genome size measurements had a relatively high standard error, and the superscaffold lengths fall within this error margin. The fact that we identified two superscaffolds as putative X chromosomes supports the hypothesis that X chromosomes are highly divergent in spiders (Kořínková & Král, 2013)..

In addition to the multiple X chromosomes, the karyotype of two distantly related entelegyne families, Agelenidae and Lycosidae (Král, 2007; Král *et al*., 2011; Sember *et al*., 2020) contains a specific XY sex chromosome pair that associates at the centromeres with the multiple X chromosomes during male meiosis (Král, 2007). This pair has been termed “cryptic sex chromosome pair” (CSCP) (Sember *et al*., 2020) since the chromosomes are homomorphic (i.e., exhibit no morphological differentiation). Available data suggest that the chromosomes forming the pair are mostly homologous (Sember et al. 2020). The CSCP can only be detected under optimal conditions, as their meiotic association with the X chromosomes is fragile. Due to its discovery in some entelegyne lineages, as well as in more basally branching spider groups, it has been hypothesized that the CSCP represents ancestral spider sex chromosomes (Král, 2007; Král *et al*., 2011, 2013; Sember *et al*., 2020). Whether or not *A. bruennichi* possesses the CSCP is beyond the scope of this study. However, if it does exist in *A. bruennichi*, the male Y and X sequences of the CSCP are very similar and the Y likely maps to the X in the genome assembly. The X_1_X_2_0 system of entelegyne spiders is thought to have arisen by X chromosome fission (e.g., (Pätau, 1948), nondisjunction of the X chromosome of an X0 system (e.g., (Postiglioni & Brum-Zorrilla, 1981), or nondisjunction of the X chromosome of the CSCP (Král, 2007). The lack of sequence similarity of X_1_ and X_2_ chromosomes found in *A. bruennichi* could reflect their origin by X chromosome fission. Alternatively, lack of sequence similarity could reflect a homologous origin via early duplication of the X chromosome in an X0 system (or early duplications of the X chromosome of CSCP) followed by a long period of differentiation of X_1_ and X_2_ chromosomes by gene mutations and chromosomal rearrangements. Recent analyses of karyotype evolution of araneomorph spiders suggest a considerable age of X_1_ and X_2_ chromosomes of the entelegyne X_1_X_2_0 system (Král *et al*., 2006; Ávila Herrera *et al*., 2021). Differentiation of X chromosome copies could be accelerated by the inactivation of X chromosome bivalents during meiosis in spider females (Král, 2007).

Interestingly, superscaffold 9 (i.e., chromosome X_1_) contains Hox cluster “A” in the *A. bruennichi* genome assembly (Sheffer *et al*., 2021). Hox genes are a set of highly conserved genes that regulate the body plan organization of animals (Pearson, Lemons, & McGinnis, 2005). This finding should be tested with fluorescence *in situ* hybridization, and investigated in other spiders, when chromosome-level assemblies and sequences for X chromosomes become available for more species. The localization of the cluster has potential implications of dosage differences in Hox genes for males and females, and the suspected compensation mechanisms should be investigated further.

Our measurements of genome size and GC content roughly matched data from the published genome assembly (Sheffer *et al*., 2021). The female measurement of 2C=4,079.809 Mbp ±256.265 corresponds to a haploid (1C) genome size of 2,039.904 Mbp ±128.132 (this study). The genome assembly has a size of 1,670.286 Mbp (Sheffer *et al*., 2021). Likewise, the average chromosome size for females was 156.916 Mbp based on flow cytometry (this study), and 126.475 Mbp in the genome assembly (Sheffer *et al*., 2021). The process of genome assembly may collapse repetitive regions, and usually results in a smaller assembly size than measured via flow cytometry (see Pflug *et al*., 2020). The GC content in flow cytometry measurements was 35.5% (this study), while the genome assembly has a GC content of 29.3%. Reduced GC content is a common trait in spiders, and the estimates reported here fall within the range reported for entelegyne spiders (27.2-35%, see Krehenwinkel *et al*., 2019; Sheffer *et al*., 2021), and indeed all other investigated spider species (Král *et al*., 2019). The GC content of females (X_1_X_1_X_2_X_2_) and males (X_1_X_2_0) was nearly identical, which indicates no substantial difference in base ratio between the X chromosomes and the autosomal set. *A. bruennichi* exhibits a larger genome than the two other studied *Argiope* species established using Feulgen densitometry, *A. aurantia* (1,584 Mbp) and *A. trifasciata* (1,653 Mbp) (Gregory & Shorthouse, 2003), but falls into the range of 1C reported for the majority of araneids. Excluding the large genome of *Hyposinga* sp. (4,000 Mbp), araneids exhibit a low diversity of genome sizes (1,438 – 2,680 Mbp) (Gregory & Shorthouse, 2003).

The DNA abundance values obtained by RT-qPCR in males for loci on superscaffold 9 show the expected pattern of approximately half abundance relative to autosomal markers and can thus be reliably used for sexing spiders. The values obtained for ABS-10C, a locus on superscaffold 10, and thus putatively sex-linked, on the other hand, resemble values obtained for autosomal loci. Given the compelling evidence from the coverage comparison that supports superscaffold 10 as being one of the two X chromosomes, we are currently unsure what might explain this phenomenon. We verified the designed primers *in silico* and found them to be specific to the region we intended to amplify. Further, the coverage for this region was neither suspiciously low nor high. However, an interpretation of the RT-qPCR results without *a priori* expectation would let us think the amplified region is placed on an autosome. While the results obtained with superscaffold 9 are robust, new primers should be designed for other fragments of superscaffold 10. The method presented here is applicable to very small DNA quantities and allowed us to identify the sex of early instar spiderlings, which are otherwise nearly impossible to sex.

Having identified the sex chromosomes in the genome assembly, future work can investigate sex chromosome evolution in *A. bruennichi*. In species in which males have one copy of each sex chromosome, as in *A. bruennichi*, the sex chromosomes are expected to evolve faster at the molecular level, relative to autosomes. This is due to several factors, such as fewer recombination events, weaker purifying selection or higher positive selection on recessive alleles relative to autosomes. These processes may lead to the acquisition of specific gene content on X chromosomes and to an X chromosome to autosome divergence (Charlesworth, Coyne, & Barton, 1987; Ellegren, 2009). A comparison of gene content between X chromosomes within and among species from diverse clades of spiders (entelegynes, haplogynes but also basally branching groups), would deliver exciting and novel insights into sex chromosome evolution and eventually of the evolution of sexual differentiation in these arachnids (Rice, 1984).

## Conclusions

We determined the *Argiope bruennichi* karyotype, genome size and GC content, which is consistent with the existing genome assembly. By comparing sequencing coverage of males and females, we were able to identify which of the superscaffolds in the genome assembly represent the multiple X chromosomes. This result enabled us to develop the first RT-qPCR markers allowing for a molecular approach to sexing spiders early during their development. Furthermore, having knowledge on the identity of the sex chromosomes in the genome assembly will allow studies on sex determination, sex allocation, sex-specific dispersal and growth, and sex-dependent physiological and developmental trajectories. Extending the taxonomic scope will allow comparisons of X chromosomes’ gene content within genomes and among species and will deliver invaluable information on the evolution of sex chromosome systems formed by X chromosomes.

## Data Availability

Sequencing reads from the male and female specimens are available on NCBI’s Short Read Archive, under BioProject PRJNA629526. Raw data for the flow cytometry data are available in the supplementary material, as is the information on RT-qPCR primers and the Ct values from the RT-qPCR experiments.

## Acknowledgements

We would like to thank Henrik Krehenwinkel (Trier, Germany) and Stefan Prost (Vienna, Austria) for discussions and brainstorming on this project early on. We also thank Lars Jensen (Greifswald, Germany) for his input on library preparation and sequencing and Douglas Araujo (Campo Grande, Brazil) for access to the paper on the karyotype of *Argiope luzona*. We are grateful to Shou-Wang Lin (Greifswald, Germany), who provided a translation of the original *Argiope bruennichi* karyotype paper, which started us on this path to generate new data. We thank the Deutsche Forschungsgemeinschaft (DFG) for the funding of this study as part of the Research Training Group 2010 RESPONSE (GRK 2010) awarded to GU. The cytogenetic part of the study was supported by the Grant Agency of Charles University (project 1000119) and the Ministry of Education, Youth, and Sports of the Czech Republic (project LTAUSA 19142). Lastly, we sincerely thank three anonymous reviewers for comments that substantially improved our manuscript.

## Author contributions

Other than the first author position, the authors are listed in alphabetical order by last name, not relative to contributions or institutional hierarchy. MMS, MC, GU, JK and MF conceived of the study. MMS, MF, and GU collected spiders. MMS and CJ isolated DNA, prepared libraries for the whole-genome sequencing and performed the sequencing, with input and infrastructure from GU and AWK. MMS, KH, MG, and MC selected loci, designed primers, and performed the RT-qPCR analysis. MK and MF performed the karyological analysis of chromosomes, with input and infrastructure from JK. EL performed the flow cytometry measurements of genome size and GC content. All authors contributed to manuscript writing and revision.

