## Supplementary for "Identification of sex chromosomes using genomic and cytogenetic methods in a range-expanding spider, *Argiope bruennichi* (Araneae: Araneidae)"

**Table S1:** Primary data of flow cytometry measurements. FI - fluorescence intensity, the mean value of a standard’s (*P. sativum*) peak or sample’s (*A. bruennichi*) peak for certain fluorochromes and their ratio. CV - coefficient of variability in (%) of each peak is given in the next column. Dye factor (DAPI) estimated as ratio of FI ratios (base specific fluorochrome / nonselective fluorochrome). Dye factor is the parameter used for calculation of base content, for details see Šmarda *et al.* (2008) and references therein.

|  | **Propidium Iodide** | | | | | **DAPI** | | | | |  |
| --- | --- | --- | --- | --- | --- | --- | --- | --- | --- | --- | --- |
| **Sex** | **FI Standard** | **CV** | **FI Sample** | **CV** | **FI sample/ FI standard** | **FI Standard** | **CV** | **FI Sample** | **CV** | **FI sample/**  **FI standard** | **Dye factor (DAPI)** |
| female | 451.38 | 1.84 | 225.21 | 4.05 | 0.4989 | 294.15 | 1.74 | 207.74 | 3.33 | 0.7062 | 1.4155 |
|  | 443.74 | 1.37 | 219.29 | 4.36 | 0.4942 | 296.26 | 1.75 | 212.14 | 2.96 | 0.7161 | 1.4490 |
|  | 410.96 | 2.22 | 206.13 | 4.87 | 0.5016 | 294.02 | 1.86 | 208.23 | 2.94 | 0.7082 | 1.4120 |
|  | 447.2 | 2.55 | 221.19 | 4.12 | 0.4946 | 302.57 | 1.61 | 215.7 | 2.51 | 0.7129 | 1.4413 |
|  | 468.57 | 2.89 | 237.5 | 4.58 | 0.5069 | 298.18 | 1.5 | 212.75 | 2.02 | 0.7135 | 1.4077 |
|  | 430.33 | 3.17 | 218.84 | 6.84 | 0.5085 | 299.98 | 1.8 | 214.04 | 2.81 | 0.7135 | 1.4031 |
|  | 412.18 | 2.19 | 228.07 | 9.91 | 0.5533 | 311.77 | 1.67 | 220.94 | 3.21 | 0.7087 | 1.2807 |
|  | 401.71 | 2.285 | 228.74 | 6.32 | 0.5694 | 313.43 | 1.76 | 222.96 | 3.14 | 0.7114 | 1.2493 |
|  | 403.76 | 2.12 | 232.85 | 5.58 | 0.5767 | 312.43 | 1.92 | 222.68 | 3.1 | 0.7127 | 1.2359 |
|  | 420.8 | 1.96 | 209.91 | 4.77 | 0.4988 | 316.22 | 1.58 | 222.62 | 2.83 | 0.7040 | 1.4113 |
| male | 473.09 | 2.48 | 210.87 | 4.48 | 0.4457 | 302.02 | 2.55 | 195.19 | 3.55 | 0.6463 | 1.4499 |
|  | 493.52 | 2.36 | 228.02 | 3.5 | 0.4620 | 303.85 | 1.58 | 196.51 | 2.67 | 0.6467 | 1.3998 |
|  | 432.38 | 1.97 | 223.47 | 8.68 | 0.5168 | 296 | 2.19 | 191.52 | 3.36 | 0.6470 | 1.2519 |
|  | 443.04 | 2.75 | 216.14 | 11.01 | 0.4879 | 297.22 | 1.93 | 190.07 | 3.44 | 0.6395 | 1.3108 |
|  | 463.8 | 2.97 | 237.37 | 5.72 | 0.5118 | 298.34 | 1.66 | 194.52 | 2.97 | 0.6520 | 1.2740 |
|  | 416.72 | 1.87 | 217.9 | 3.29 | 0.5229 | 324.04 | 1.96 | 205.67 | 5.75 | 0.6347 | 1.2138 |
|  | 461.34 | 6.17 | 235.43 | 4.77 | 0.5103 | 315.27 | 1.9 | 201.56 | 3.85 | 0.6393 | 1.2528 |
|  | 398.67 | 1.71 | 206.25 | 5.43 | 0.5173 | 310.63 | 2.13 | 203.1 | 4.21 | 0.6538 | 1.2638 |
|  | 400.73 | 1.97 | 208.36 | 5.62 | 0.5200 | 309.77 | 1.89 | 198.96 | 5.16 | 0.6423 | 1.2353 |
|  | 388.22 | 2.75 | 196.37 | 6.28 | 0.5058 | 312.39 | 2.19 | 201.58 | 4.02 | 0.6453 | 1.2757 |

**Table S2**: DNA quantities obtained prior to RT-qPCR or conventional PCR (“conv. PCR”), and triplicate Ct values for each locus in each individual from RT-qPCR. A minus sign (-) indicates no or missing data, i.e. if a marker was not run or one replicate was excluded due to high deviation. The extraction column lists the extraction method used (Promega kit (“promega”) or basic principles extraction (“basic”)).

| **Sex** | **Individual ID** | **Nanodrop (ng µL^-1^)** | **Qubit (ng µL^-1^)** | **Extraction** | **Use** | **Locus** | **Ct1** | **Ct2** | **Ct3** |
| --- | --- | --- | --- | --- | --- | --- | --- | --- | --- |
| Female | Ab6 | 58.4 | 14.6 | promega | RT-qPCR | ABA-5 | 24.48 | 26.96 | 24.48 |
|  |  |  |  |  |  | ABA-7 | 22.75 | 22.77 | 22.80 |
|  |  |  |  |  |  | ABS-9.3 | 29.76 | 29.90 | 29.95 |
|  |  |  |  |  |  | ABS-9.4 | 25.74 | 25.77 | 25.76 |
|  |  |  |  |  |  | ABS-9.5 | 25.31 | 25.36 | 25.37 |
|  |  |  |  |  |  | ABS-10C | 21.66 | 21.64 | 21.65 |
|  | Ab7 | 39.8 | 13.7 | promega | RT-qPCR | ABA-5 | 21.47 | 21.43 | 21.41 |
|  |  |  |  |  |  | ABA-7 | 27.49 | 27.44 | 27.55 |
|  |  |  |  |  |  | ABS-9.3 | 27.06 | 27.31 | 27.23 |
|  |  |  |  |  |  | ABS-9.4 | 25.72 | 25.64 | 25.98 |
|  |  |  |  |  |  | ABS-9.5 | 26.63 | 26.71 | 26.74 |
|  |  |  |  |  |  | ABS-10C | 21.16 | 21.06 | 21.05 |
|  | Ab2 | 103.60 | 31.6 | promega | conv. PCR | - | - | - | - |
|  | Ab3 | 78 | NA | promega | conv. PCR | - | - | - | - |
|  | AB_F1 | 267.0 | 26.6 | basic | RT-qPCR | ABA-5 | 21.85 | 21.95 | 21.92 |
|  |  |  |  |  |  | ABA-7 | 22.36 | 22.66 | 22.65 |
|  |  |  |  |  |  | ABS-9.3 | 22.63 | 22.60 | 22.81 |
|  |  |  |  |  |  | ABS-9.4 | 23.10 | 23.10 | 23.15 |
|  |  |  |  |  |  | ABS-9.5 | 22.64 | 22.67 | 22.81 |
|  |  |  |  |  |  | ABS-10C | 19.41 | 19.33 | 19.29 |
|  | AB_F2 | 436.4 | 75.4 | basic | RT-qPCR | ABA-5 | 22.67 | 22.64 | 22.95 |
|  |  |  |  |  |  | ABA-7 | 22.98 | 23.13 | 23.20 |
|  |  |  |  |  |  | ABS-9.3 | 23.18 | 23.22 | 23.21 |
|  |  |  |  |  |  | ABS-9.4 | 23.61 | 23.65 | 23.70 |
|  |  |  |  |  |  | ABS-9.5 | 23.12 | 23.25 | 23.94 |
|  |  |  |  |  |  | ABS-10C | 19.90 | 19.71 | 19.78 |
|  | AB_F3 | 250.2 | 94.2 |  | RT-qPCR | ABA-5 | 24.67 | 24.68 | 24.71 |
|  |  |  |  |  |  | ABA-7 | 25.04 | 25.07 | 25.05 |
|  |  |  |  |  |  | ABS-9.3 | 25.12 | 25.11 | 25.18 |
|  |  |  |  |  |  | ABS-9.4 | 25.65 | 25.67 | 25.75 |
|  |  |  |  |  |  | ABS-9.5 | 24.91 | 24.96 | 24.97 |
|  |  |  |  |  |  | ABS-10C | 22.24 | 22.22 | 23.24 |
|  | AB_F4 | 154.9 | 50.4 | basic | RT-qPCR | ABA-5 | 24.21 | 24.27 | 24.43 |
|  |  |  |  |  |  | ABA-7 | 24.53 | 24.67 | 24.63 |
|  |  |  |  |  |  | ABS-9.3 | 24.72 | 24.76 | 24.79 |
|  |  |  |  |  |  | ABS-9.4 | 25.21 | 25.20 | 25.12 |
|  |  |  |  |  |  | ABS-9.5 | 24.47 | 24.54 | 24.85 |
|  |  |  |  |  |  | ABS-10C | 21.41 | - | 21.56 |
| Male | Ab26 | 11.1 | 5.17 | promega | RT-qPCR | ABA-5 | 25.00 | 24.13 | 24.12 |
|  |  |  |  |  |  | ABA-7 | 25.03 | 22.43 | 22.49 |
|  |  |  |  |  |  | ABS-9.3 | 30.66 | 30.60 | 30.54 |
|  |  |  |  |  |  | ABS-9.4 | 27.01 | 26.46 | 26.51 |
|  |  |  |  |  |  | ABS-9.5 | 26.07 | 25.92 | 25.97 |
|  |  |  |  |  |  | ABS-10C | 21.97 | 20.97 | 20.95 |
|  | Ab27 | 12.4 | 8.53 | promega | RT-qPCR | ABA-5 | 22.41 | 22.14 | 22.57 |
|  |  |  |  |  |  | ABA-7 | 27.96 | 27.82 | 27.8 |
|  |  |  |  |  |  | ABS-9.3 | 29.98 | 28.49 | 28.46 |
|  |  |  |  |  |  | ABS-9.4 | 27.17 | 27.12 | 26.99 |
|  |  |  |  |  |  | ABS-9.5 | 28.96 | 27.96 | 28.32 |
|  |  |  |  |  |  | ABS-10C | 21.94 | 21.85 | 21.72 |
|  | AB_M1 | 80.17 | 24.4 | basic | RT-qPCR | ABA-5 | 22.66 | 22.65 | 22.66 |
|  |  |  |  |  |  | ABA-7 | 22.62 | 22.70 | 22.75 |
|  |  |  |  |  |  | ABS-9.3 | 23.93 | 23.77 | 23.70 |
|  |  |  |  |  |  | ABS-9.4 | 24.11 | 24.03 | 24.01 |
|  |  |  |  |  |  | ABS-9.5 | 23.77 | 23.68 | 23.73 |
|  |  |  |  |  |  | ABS-10C | 19.46 | 19.43 | 19.38 |
|  | AB_M2 | 107.8 | 39.0 | basic | RT-qPCR | ABA-5 | 23.00 | 23.22 | 23.02 |
|  |  |  |  |  |  | ABA-7 | 22.91 | 22.94 | 23.09 |
|  |  |  |  |  |  | ABS-9.3 | 24.14 | 24.22 | 24.18 |
|  |  |  |  |  |  | ABS-9.4 | 24.63 | 24.57 | 24.51 |
|  |  |  |  |  |  | ABS-9.5 | 24.28 | 24.29 | 24.37 |
|  |  |  |  |  |  | ABS-10C | 19.96 | 19.52 | 19.83 |
|  | AB_M3 | 25.30 | 12.7 | basic | RT-qPCR | ABA-5 | 24.62 | 24.48 | 24.88 |
|  |  |  |  |  |  | ABA-7 | 24.63 | 24.77 | 24.74 |
|  |  |  |  |  |  | ABS-9.3 | 25.98 | 26.00 | 25.97 |
|  |  |  |  |  |  | ABS-9.4 | 26.45 | 26.53 | 26.37 |
|  |  |  |  |  |  | ABS-9.5 | 25.82 | 25.80 | 25.72 |
|  |  |  |  |  |  | ABS-10C | 22.36 | 22.42 | 22.38 |
|  | AB_M4 | 31.75 | 10.5 | basic | RT-qPCR | ABA-5 | 24.14 | 24.33 | 24.26 |
|  |  |  |  |  |  | ABA-7 | 24.71 | 24.78 | 24.73 |
|  |  |  |  |  |  | ABS-9.3 | 25.86 | 25.45 | 25.92 |
|  |  |  |  |  |  | ABS-9.4 | 26.25 | 26.34 | 26.28 |
|  |  |  |  |  |  | ABS-9.5 | 25.58 | 25.60 | 25.95 |
|  |  |  |  |  |  | ABS-10C | 21.90 | 22.02 | 21.97 |
| 1^st^ instar spiderlings | AB_S1 | 35.76 | 42.8 | basic | RT-qPCR | ABA-7 | 24.59 | 24.56 | 24.73 |
|  |  |  |  |  |  | ABS-9.3 | 25.72 | 25.39 | 25.51 |
|  |  |  |  |  |  | ABS-9.4 | 25.88 | 25.77 | 25.79 |
|  |  |  |  |  |  | ABS-9.5 | 25.71 | 25.68 | 25.60 |
|  | AB_S2 | 23.24 | 22.6 | basic | RT-qPCR | ABA-7 | 24.12 | 24.05 | 24.18 |
|  |  |  |  |  |  | ABS-9.3 | 25.25 | 24.92 | 24.97 |
|  |  |  |  |  |  | ABS-9.4 | 25.15 | 25.17 | 25.12 |
|  |  |  |  |  |  | ABS-9.5 | 24.97 | 24.90 | 24.99 |
|  | AB_S3 | 32.07 | 18.0 | basic | RT-qPCR | ABA-7 | 23.90 | 24.02 | 24.02 |
|  |  |  |  |  |  | ABS-9.3 | 23.80 | 23.84 | 23.78 |
|  |  |  |  |  |  | ABS-9.4 | 24.04 | 24.10 | 24.20 |
|  |  |  |  |  |  | ABS-9.5 | 23.74 | 23.90 | 23.88 |
|  | AB_S4 | 25.89 | 27.0 | basic | RT-qPCR | ABA-7 | 24.18 | 24.24 | 24.56 |
|  |  |  |  |  |  | ABS-9.3 | 24.14 | 24.10 | 24.11 |
|  |  |  |  |  |  | ABS-9.4 | 24.33 | 24.44 | 24.45 |
|  |  |  |  |  |  | ABS-9.5 | 24.11 | 24.28 | 24.25 |
|  | AB_S5 | 27.50 | 30.4 | basic | RT-qPCR | ABA-7 | 24.47 | 24.54 | 24.55 |
|  |  |  |  |  |  | ABS-9.3 | 25.49 | 25.39 | 25.48 |
|  |  |  |  |  |  | ABS-9.4 | 25.92 | 26.12 | 25.99 |
|  |  |  |  |  |  | ABS-9.5 | 25.81 | 25.67 | 25.63 |
|  | AB_S6 | 31.16 | 28.8 | basic | RT-qPCR | ABA-7 | 23.92 | 23.96 | 24.07 |
|  |  |  |  |  |  | ABS-9.3 | 24.29 | 23.94 | 23.88 |
|  |  |  |  |  |  | ABS-9.4 | 24.24 | 24.19 | 24.09 |
|  |  |  |  |  |  | ABS-9.5 | 23.99 | 23.90 | 23.98 |
|  | AB_S7 | 30.46 | 23.0 | basic | RT-qPCR | ABA-7 | 23.91 | 23.93 | 23.89 |
|  |  |  |  |  |  | ABS-9.3 | 24.78 | 24.76 | 24.77 |
|  |  |  |  |  |  | ABS-9.4 | 24.98 | 24.97 | 25.14 |
|  |  |  |  |  |  | ABS-9.5 | 24.80 | 24.87 | 24.83 |
|  | AB_S8 | 43.58 | 30.0 | basic | RT-qPCR | ABA-7 | 24.20 | 24.20 | 24.22 |
|  |  |  |  |  |  | ABS-9.3 | 24.07 | 24.13 | 24.12 |
|  |  |  |  |  |  | ABS-9.4 | 24.36 | 24.34 | 24.43 |
|  |  |  |  |  |  | ABS-9.5 | 24.10 | 24.13 | 24.15 |
|  | AB_S9 | 26.27 | 18.9 | basic | RT-qPCR | ABA-7 | 23.73 | 23.91 | 23.75 |
|  |  |  |  |  |  | ABS-9.3 | 24.61 | 24.66 | 24.68 |
|  |  |  |  |  |  | ABS-9.4 | 24.91 | 24.87 | 24.90 |
|  |  |  |  |  |  | ABS-9.5 | 24.56 | 24.67 | 24.70 |
|  | AB_S10 | 21.03 | 19.5 | basic | RT-qPCR | ABA-7 | 25.10 | 25.15 | 25.31 |
|  |  |  |  |  |  | ABS-9.3 | 25.29 | 25.32 | 25.44 |
|  |  |  |  |  |  | ABS-9.4 | 25.74 | 25.77 | 25.85 |
|  |  |  |  |  |  | ABS-9.5 | 25.13 | 25.22 | 25.19 |

**Table S3**: Primers used for quantification in the RT-qPCR approach. ABA-5 and ABA-7 are autosomal loci, while ABS-10C, ABS-9.3, ABS-9.4 and ABS-9.5 should be sex-linked. Primers marked with an asterisk (*) were used to sex spiderlings. The two efficiencies provided in the final column represent different efficiencies in different labs (Hamburg/Greifswald).

| **Locus** | **Scaff** | **Forward (“fw”)** | **Reverse (“rv”)** | **Product Size** [bp] | **Temp fw** [°C] | **Temp rv** [°C] | **Efficiency** [%] |
| --- | --- | --- | --- | --- | --- | --- | --- |
| ABA-5 | 5 | caagccgactgatagagatccc | gaccgaacattattaaccgccg | 151 | 60.03 | 60.03 | 108/147 |
| ABA-7* | 7 | ACGAGATGTACTGACGGCATAG | TTCCGCTGGTTCTTTTGCATTC | 171 | 59.71 | 60.29 | 107/102 |
| ABS-9.3* | 9 | GTGGCACTTTTAGCTGGAACTG | ttcttacCCATCAAAGGAGGCC | 176 | 60.03 | 60.03 | 83/108 |
| ABS-9.4* | 9 | agaggagagttggttggctttc | ccacttcgcgcatctgaataag | 188 | 60.22 | 60.03 | 109/105 |
| ABS-9.5* | 9 | cttcttattttggcgggctagc | catcaagcagtgttgtggaagg | 158 | 59.97 | 60.03 | 96/97 |
| ABS-10C | 10 | ATTCATGTTCGCCGCATTAGTG | GCAGTCCTCAAATGTCCAAACC | 191 | 59.97 | 60.03 | 97/113 |

**Table S4:** Fold-change values of sex-linked loci in spiderlings. Values are rounded to two decimal places.

| **ID** | **Fold-change**  **ABS-9.3** | **Fold-change**  **ABS-9.4** | **Fold-change**  **ABS-9.5** | **Mean**  **fold-change** | **Probable sex** |
| --- | --- | --- | --- | --- | --- |
| S1 | 0.45 | 0.48 | 0.48 | 0.47 | male |
| S2 | 0.45 | 0.54 | 0.54 | 0.51 | male |
| S3 | 1.04 | 1.05 | 1.04 | 1.05 | female |
| S4 | 1.05 | 1.08 | 1.03 | 1.06 | female |
| S5 | 0.51 | 0.48 | 0.59 | 0.53 | male |
| S6 | 0.79 | 0.97 | 0.89 | 0.88 | female |
| S7 | 0.44 | 0.49 | 0.47 | 0.47 | male |
| S8 | 0.89 | 0.98 | 0.94 | 0.93 | female |
| S9 | 0.44 | 0.50 | 0.49 | 0.48 | male |
| S10 | 0.94 | 1.03 | 0.94 | 0.97 | female |
